# Maturation and conformational switching of a de novo designed phase-separating polypeptide

**DOI:** 10.1101/2024.01.08.574599

**Authors:** Alexander T. Hilditch, Andrey Romanyuk, Lorna R. Hodgson, Judith Mantell, Chris R. Neal, Paul Verkade, Richard Obexer, Louise C. Serpell, Jennifer J. McManus, Derek N. Woolfson

**Affiliations:** School of Chemistry, University of Bristol, Cantock’s Close, Bristol, BS8 1TS, UK; Max Planck-Bristol Centre for Minimal Biology, University of Bristol, Cantock’s Close, Bristol, BS8 1TS, UK; Wolfson Bioimaging Facility, University of Bristol, Biomedical Sciences Building, Bristol, BS8 1TD, UK; School of Biochemistry, University of Bristol, Biomedical Sciences Building, Bristol, BS8 1TD, UK; Bristol BioDesign Institute, School of Chemistry, University of Bristol, Cantock’s Close, Bristol, BS8 1TS, UK; Department of Chemistry, Manchester Institute of Biotechnology, University of Manchester, Princess Street, Manchester M1 7DN, UK; School of Life Sciences, University of Sussex, Falmer, Brighton, JMS 3B17, UK; HH Wills Physics Laboratory, School of Physics, University of Bristol, Tyndall Avenue, Bristol, BS8 1TL, UK

**Keywords:** Liquid-liquid phase separation, membraneless organelles, protein-protein interactions, protein design, protein fibers, amyloid

## Abstract

Cellular compartments formed by biomolecular condensation are a widespread feature of cell biology. These organelle-like assemblies compartmentalize macromolecules dynamically within the crowded intracellular environment. However, the intermolecular interactions that produce condensed droplets may also create arrested states and potentially pathological assemblies such as fibers, aggregates, and gels, through droplet maturation. Protein liquid-liquid phase separation is a metastable process, so maturation may be an intrinsic property of phase-separating proteins, where nucleation of different phases or states arise in supersaturated condensates. Here, we describe the formation of both phase-separated droplets and proteinaceous fibers driven by a de novo designed polypeptide. We characterize the formation of supramolecular fibers in vitro and in bacterial cells. We show that client proteins can be targeted to the fibers in cells using the droplet-forming construct. Finally, we explore the interplay between phase separation and fiber formation of the de novo polypeptide, showing that the droplets mature with a post-translational switch to largely β conformations, analogous to models of pathological phase separation.

## MAIN TEXT

Membraneless organelles (MLOs) formed by biomolecular condensation are now recognized as a widespread phenomenon in cell biology.^1^ Macromolecular condensation is implicated in many cellular processes, from chromatin organization, cell signaling, and genetic regulation, to microtubule organization, cell division, and cellular stress responses.^2-5^ Nonetheless, there are still significant gaps in our knowledge of how phase-separating proteins are organized and controlled at a molecular level.^6^ Recent advances in understanding the molecular driving forces of protein phase separation have come from the bottom-up design of artificial biomolecular condensates.^7^ These offer the potential to create synthetic MLOs, orthogonal to and isolated from endogenous cellular processes. While phase separation is being explored by synthetic biologists for its potential to augment biology, the same processes are being interrogated clinically for their potential roles in disease and degenerative processes, such as frontotemporal dementia (FTD), and amyotrophic lateral sclerosis (ALS).^8^

In protein phase separation, weak attractive intermolecular interactions produce de-mixed droplets in which macromolecules are enriched, but still exchange and diffuse quickly.^9^ However, the same interactions that drive self-assembly of dynamic droplets can also lead to arrested proteinaceous assemblies. Indeed, the latter can nucleate from phase-separated droplets.^10^ This secondary process which nucleates the formation of a new protein state has been termed the maturation or molecular ageing of protein condensates.^11^ In some cases, droplet maturation is an essential part of biological function.^12^ However, droplet maturation has been implicated in the formation of pathological proteinaceous assemblies, including amyloid fibers and aggregates.^13,14^ Moreover, several ALS-related mutations in phase-separating proteins have been observed to accelerate the transition from dynamic to arrested states. ^10,15,16^

The mechanisms that lead to droplet maturation are relatively poorly understood. Some maturation events have been linked to specific sequence elements, or changes in the cellular environment.^17^ Understanding how droplet maturation occurs could lead to more effective treatment of amyloid pathologies.^18^ Previously, we reported a *de novo* polypeptide, HERD-2.2, for liquid-liquid phase separation (LLPS, Table S1).^19^ Here, we describe the interplay between phase separation and fiber formation of this synthetic polypeptide. HERD-2.2 was designed to function as a genetic fusion to client proteins, such as fluorescent reporters or enzymes. For instance, a fusion to the green fluorescent protein (GFP) mEmerald, denoted HERD-2.2–GFP, phase separates as de-mixed droplets *in vitro* and in *Escherichia coli* (*E. coli*) (Figures 1A & S1). Here we describe the phase behavior of the isolated HERD-2.2 module in isolation.

**Figure 1.**
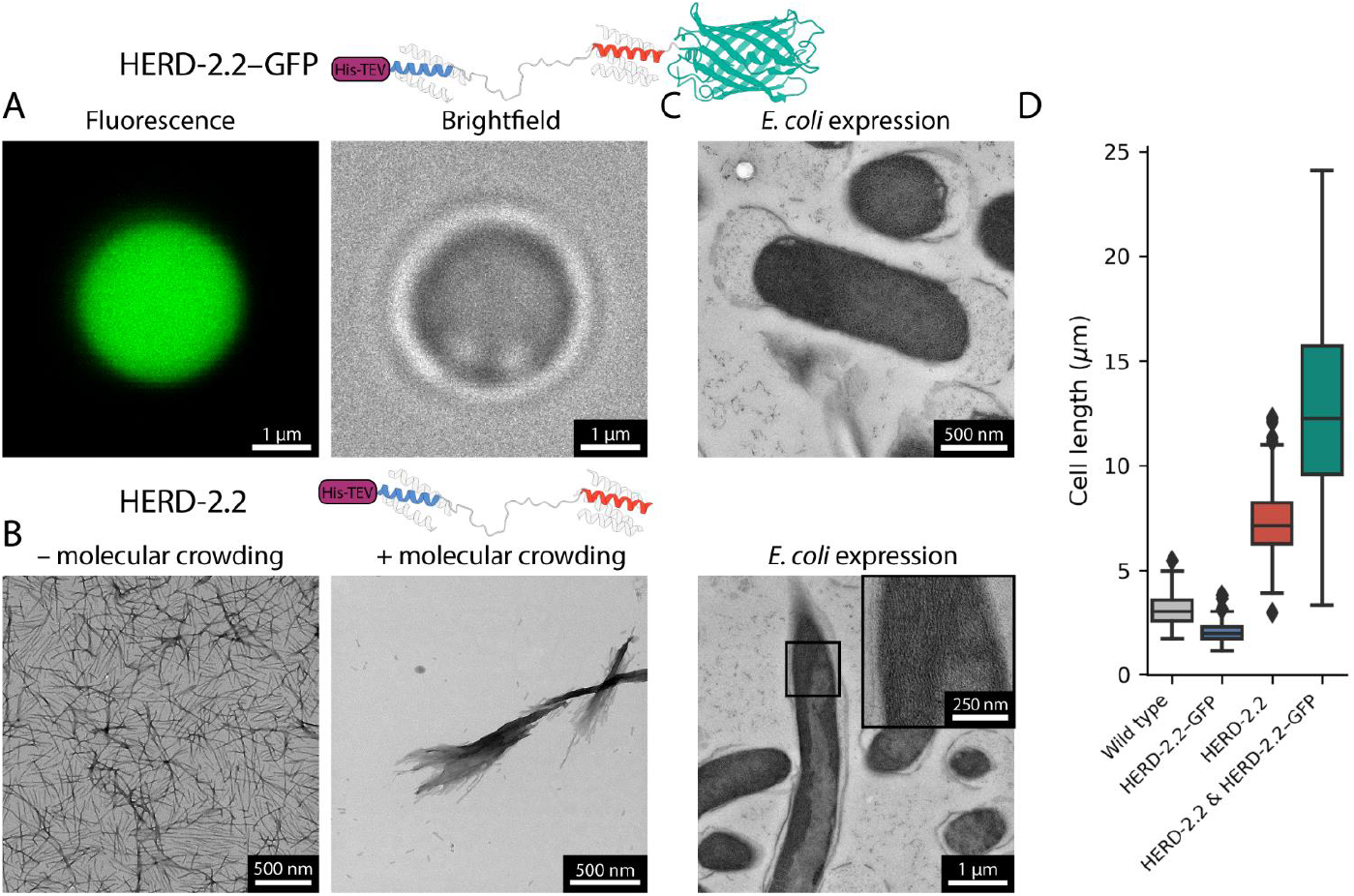
HERD-2.2 forms supramolecular fibers *in vitro* and in *E. coli*. (A) Confocal microscopy images of HERD-2.2–GFP. Conditions: 110 μM HERD-2.2–GFP, 10% PEG 3350, 125 mM NaCl, 50 mM Tris pH 7.5. (B) Negative stain TEM images of HERD-2.2 without molecular crowding (left; 25 μM HERD-2.2, 50 mM Tris pH 7.5) and with (right; 25 μM HERD-2.2, 10% PEG 3350, 125 mM NaCl, 50 mM Tris pH 7.5). (C) TEM sections of *E. coli* expressing HERD-2.2–GFP and HERD-2.2. (D) Cell length of WT *E. coli* (n=106), *E. coli* expressing HERD-2.2--GFP (n=100), HERD-2.2 (n=107), and HERD-2.2--GFP and HERD-2.2 co-expressed (n=100).

The isolated HERD-2.2 polypeptide was expressed and purified from *E. coli* using an *N*-terminal His tag, and phase separation probed *in vitro*. Instead of forming de-mixed droplets, HERD-2.2 formed supramolecular fibers (Figures 1B & S2). The HERD-2.2 fibers were examined by negative stain transmission electron microscopy (TEM). This revealed dispersed individual protein fibers in dilute solution, with a diameter of 13 nm (Figure S3). However, with the addition of molecular crowding agents to mimic the cytoplasmic environment, and as used to phase separate HERD-2.2–GFP (10% PEG 3350, 125 mM NaCl, 50 mM Tris pH 7.5), the fibers assembled further and laterally to give structures several microns in length and of variable thickness (Figure S4).

Having observed HERD-2.2 fibers *in vitro*, we looked directly in cells. Sections of *E. coli* expressing HERD-2.2 were imaged by TEM, confirming fiber formation within the cytoplasm (Figures 1C & S5). These fibers assembled laterally, analogous to those seen *in vitro* with molecular crowding agents. Cells expressing HERD-2.2–GFP did not form fibers, but the previously reported dense regions of protein at the cell poles (Figures S6 & S7). Moreover, cells expressing HERD-2.2 grew much longer than the expected length for *E. coli* with a mean length of 7.4 ± 1.8 µm, while cells expressing HERD-2.2– GFP condensates grew to lengths of 2.0 ± 0.6 µm, slightly shorter than wild type (WT) cells at 3.1 ± 0.7 µm (Figure 1D).

The formation of such large fibers within the *E. coli* cytoplasm is remarkable, and reminiscent of engineered systems for subcellular recruitment using fibrous assemblies.^20^ Therefore, we assessed the capacity for HERD-2.2 fibers to act as a recruitment scaffold, but in this case for other assemblies. To this end, HERD-2.2 and HERD-2.2–GFP were co-expressed in *E. coli*. This gave fluorescently labelled fibers, demonstrating that GFP had been recruited to the scaffold fibers (Figure S8). These cells shared the same elongated phenotype as cells expressing HERD-2.2 alone, growing to a mean length of 12.8 ± 4.5 µm (Figure 1D). To confirm recruitment of HERD-2.2–GFP to the intracellular fibers, cells co-expressing HERD-2.2 and HERD-2.2–GFP were examined by correlative light and electron microscopy (CLEM). This revealed fluorescent HERD-2.2–GFP enriched along the HERD-2.2 fibers (Figure 2A), while expression of HERD-2.2–GFP alone produced fluorescence only around the condensates at the cell poles (Figure S9). Moreover, HERD-2.2 co-expressed with both HERD-2.2–GFP and HERD-2.2– mCherry showed enrichment of both fluorescent proteins; while co-expression with non-tagged GFP and HERD-2.2–mCherry only the latter was recruited to the fibers (Figure S10).

**Figure 2.**
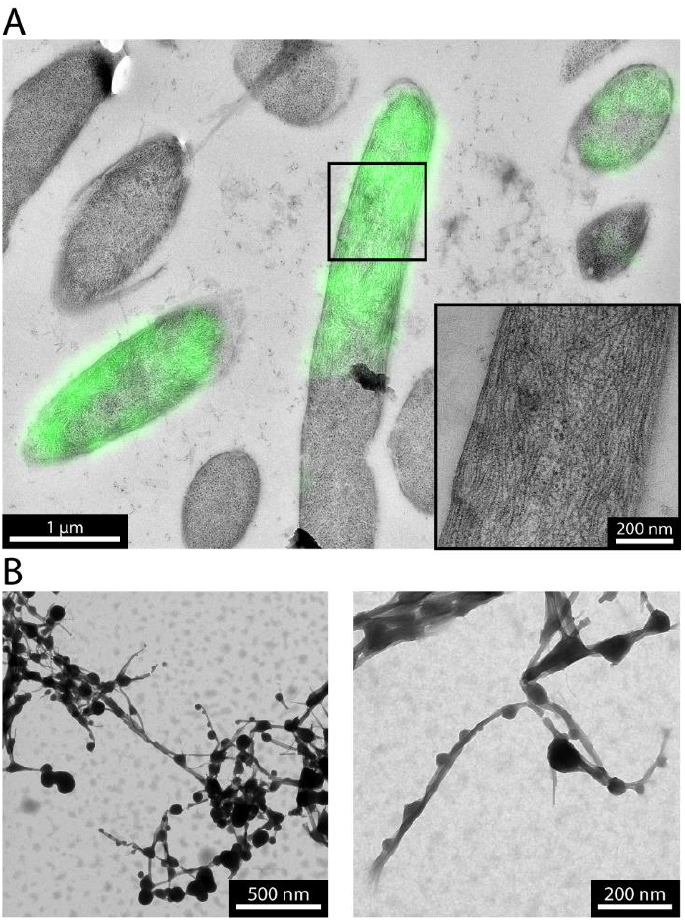
HERD-2.2 fibers labelled with droplet-forming HERD-2.2–GFP. (A) CLEM images of *E. coli* co-expressing HERD-2.2 and HERD-2.2–GFP. (B) TEM images of mixtures of HERD-2.2 and HERD-2.2–GFP. Conditions: 82 μM HERD-2.2, 55 μM HERD-2.2–GFP, 10% PEG 3350, 125 mM NaCl, 50 mM Tris pH 7.5.

Next, we examined the HERD-2.2–GFP and HERD-2.2 composite *in vitro*. Individually, these formed de-mixed liquid droplets and elongated fibers respectively. When combined, and with molecular crowding regents, fluorescently labelled fibers were formed as observed in *E. coli*. However, these cell-free experiments revealed that the fluorescent protein was not distributed evenly, but formed puncta along the fibers (Figure S11). This indicates that multi-phase assemblies of both fibrous, and droplet-like structures can be formed independently, but that they have an affinity for each other, which we suggest arises from the common HERD-2.2 component. TEM of these mixtures confirmed that both phases were formed, leading to laterally assembled elongated fibers decorated with spherical droplet-like structures along their lengths (Figures 2B, S4 & S11).

Having observed fiber assembly, we interrogated their molecular structure. Initially, we postulated that the fibers could be α helical, as helical protein-protein interactions were used to design the HERD-2.2 fusions.^19^ Therefore, we examined the structure of the fibers by circular dichroism (CD) spectroscopy. CD spectra of the fibers in solution at 5 °C showed a weak shoulder around 220 nm (Figure 3A). Deconvolution of the spectrum indicated largely unstructured protein, with some α and β contribution (Figure S12). Heating the sample from 5 – 90 °C resulted in the irreversible loss of the shoulder and the α helical contribution of the deconvolution (Figure S12), with a melting temperature of 58.0 ± 0.5 °C (Figure 3B). TEM samples before and after melting confirmed the loss of fibrous structures (Figure S13). However, cooling back to 5 °C did not induce re-folding, nor the reformation of fibers. To elucidate the structure of the fibers further, we turned to X-ray fiber diffraction. Solutions of fibers were suspended between glass capillaries and partially dried, to produce aligned fibers (Figure S14). Diffraction of these gave a strong meridional reflection at 4.72 Å, characteristic of a cross-β structure (Figures 3C & S15).^21,22^ It is possible that this β structure was a consequence of partially drying the sample under tension. Therefore, we investigated the interaction of HERD-2.2 fibers with Thioflavin T (ThT). ThT shows increased fluorescence on interaction with amyloid-like fibrils.^23^ As a control, we also incubated ThT with a *de novo* dimeric α-helical coiled-coil peptide CC-Di.^24^

**Figure 3.**
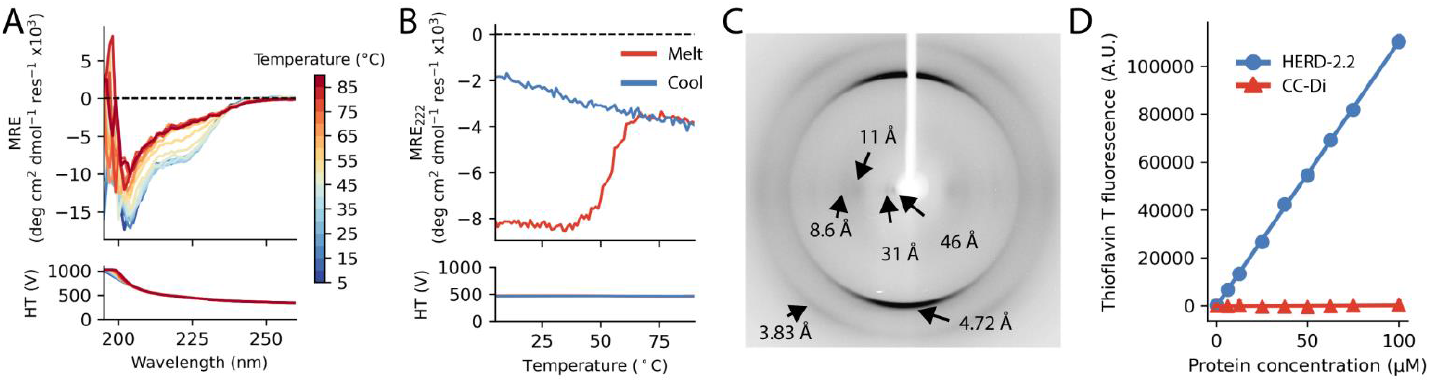
Biophysical characterization of HERD-2.2 secondary structure. (A) CD spectra of HERD-2.2 on heating from 5 – 90 °C. Conditions: 25 μM HERD-2.2, 50 mM Tris pH 7.5. (B) CD of HERD-2.2 at 222 nm on heating and cooling. (C) X-ray fiber diffraction pattern of HERD-2.2. (D) Thioflavin T fluorescence with addition of HERD-2.2 or CC-Di.

Addition of HERD-2.2 to 25 µM solutions of ThT resulted in a linear increase in ThT fluorescence proportional to the HERD-2.2 concentration, as seen for amyloid fibrils (Figure 3D).^25^ Moreover, the fold-increase in ThT fluorescence closely matched that measured for the amyloid fibrils Aβ40 and Aβ42 (approximately 3.1-fold at 8 μM protein).^25^ In contrast, addition of CC-Di gave no increase in ThT fluorescence, even with up to 100 μM peptide.

This possible amyloid-like structure was surprising, as it is not consistent with the CD spectra of the peptide constituents.^19^ Nonetheless, it suggests a structural transition to β is possible in these constructs. Moreover, β secondary structure indicates that the HERD-2.2 fibers may be similar to the amyloid-like assemblies formed by natural phase-separating proteins;^26,27^ although the fibers formed by HERD-2.2 are more thermally labile than is usually observed with amyloid-like assemblies (Figure 3B).^28,29^

The formation of both droplets and fibers by a designed protein could provide an accessible synthetic system for studying droplet maturation. To understand this change in phase behavior further, we assessed whether a HERD-2.2 system could be switched from liquid-like droplets to fibers *in situ* by post-translational processing. The key difference between these two phase behaviors is the addition of GFP *C*-terminal to the HERD-2.2 polypeptide. Therefore, to introduce a triggerable response, we inserted a thrombin cleavage site between HERD-2.2 and GFP, forming HERD-2.2-T–GFP. This protein was expressed and purified from *E. coli*. As with the parent HERD-2.2–GFP, addition of molecular crowding agents caused HERD-2.2-T–GFP to de-mix and form fluorescent droplets in solution (Figure 4A). Next, we added the protease thrombin to cleave the *C*-terminal fluorescent protein. SDS-PAGE indicated that cleavage was complete after 1 hour, and was associated with clearing of the solution, indicative of some droplets dispersing following cleavage (Figures S16 & S17). Examination of the remaining droplets by confocal microscopy indicated a marked maturation effect, with coarsening and roughening of their surfaces (Figures 4A & S18). TEM of the mature droplets confirmed the differences between these assemblies.

**Figure 4.**
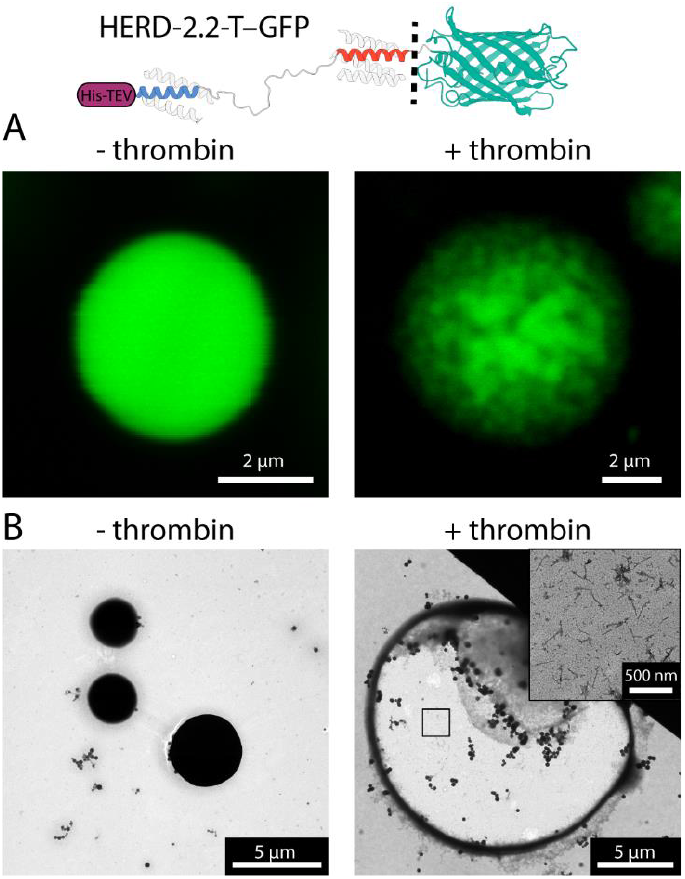
Maturation and fiber formation from HERD-2.2-T–GFP droplets. Confocal fluorescence microscopy (A) and TEM (B) of HERD-2.2-T-GFP with and without protease. Conditions: 10% PEG 3350, 125 mM NaCl, 50 mM Tris pH 7.5, 0.675 mM HERD-2.2-T–GFP with (+ thrombin) or without (– thrombin) addition of 0.06 units of thrombin protease.

In the absence of protease, HERD-2.2-T–GFP formed highly dense droplets without evident internal structure (Figures 4B & S19). Following protease cleavage however, the droplets formed hollowed-out macroscopic assemblies, with a visible proteinaceous corona. Moreover, within these assemblies rod-shaped fibers were observable, similar to those formed by HERD-2.2. Thus, the transition from de-mixed liquid to fiber had been triggered by the cleavage.

In summary, we have characterized a synthetic polypeptide, HERD-2.2, that drives the formation of both phase-separated droplets, and macromolecular fibers, which assemble both *in vitro* and in cells. Previously, we used LLPS of HERD-2.2–GFP to colocalize functional enzymes, conferring a boost to a two-enzyme pathway.^19^ Here, we have demonstrated that HERD-2.2 fibers can also act as recruitment scaffolds in cells. This presents opportunities to study localization of clients using both rigid and dynamic assemblies. Moreover, we have identified that the *de novo* polypeptide can be switched between de-mixed droplets, and mature fibrous structures, and that switching is associated with a conformational change to β secondary structure. A switch from α to β secondary structure has been reported previously in other *de novo* designed peptides.^30^ The phase-separating protein TDP-43 has also been identified to develop additional β structure during early droplet maturation events, eventually forming an amyloid-like state^31^, and phase separation has been shown to promote β structure in Tau repeats.^32^ There are indications that a similar process may occur in FUS, which forms hardened β shells during droplet ageing.^33^ This suggests that this process in the *de novo* HERD-2.2 may be recapitulating intrinsic properties of phase-separating proteins. All of TDP-43, Tau, and FUS form pathological aggregates linked to the neurodegenerative disorders Alzheimer’s, FTD, and ALS.^34-36^ The HERD-2.2 system could be of use in mimicking and following how droplet maturation occurs at the molecular level, providing insights into the nucleation and growth of pathogenic aggregates. Also, as demonstrated here, HERD-2.2 assembly in living cells makes them interesting for studying the effects of supramolecular polypeptide structures on cell biology. Droplet templating and maturation could be a viable strategy for creating microscopically structured proteinaceous materials.

## AUTHOR INFORMATION

### Author Contributions

ATH, RO, JJM, and DNW conceived the project. Confocal microscopy was performed by ATH. TEM was performed by ATH, LRH, and JM. CLEM was performed by ATH, LRH and CRN with assistance from PV. CD was performed by ATH and AR. X-ray fiber diffraction was performed by ATH and LCS. Droplet maturation experiments were performed by ATH. ATH, JJM, and DNW wrote the manuscript. All authors read and contributed to the preparation of the manuscript.

### Notes

The authors declare no competing interests.

## ACKNOWLEDGMENT

The authors are grateful to the Wolfson Bioimaging Facility, University of Bristol, for access to confocal and transmission electron microscopes, as well as their help and support. We thank Katherine Albanese for assistance with Thioflavin T measurements, and Kate Kurgan for supplying purified CC-Di peptide.

## ABBREVIATIONS

LLPS: liquid-liquid phase separation
MLO: membraneless organelle
GFP: green fluorescent protein
HERD: helical repeat domain
TEM: transmission electron microscopy
WT: wild type
CLEM: correlative light and electron microscopy
CD: circular dichroism.

