## Supporting information for "Maturation and conformational switching of a de novo designed phase-separating polypeptide"

### Contents

Materials and methods ... 3
- Materials ... 3
- Computational tools ... 3
- Protein expression and purification ... 3
- Confocal microscopy ... 3
- Transmission electron microscopy (TEM) ... 4
- Correlative light electron microscopy (CLEM) ... 4
- Circular dichroism (CD) spectroscopy ... 4
- X-ray fiber diffraction ... 5
- Thioflavin T fluorescence ... 5
- Protease cleavage and droplet maturation ... 5

Supporting information ... 6
- Supporting table S1. Protein sequences used in this study. ... 6
- Supporting figure S1. Additional confocal microscopy images of HERD-2.2–GFP droplets ... 7
- Supporting figure S2. Additional TEM images of HERD-2.2 fibers. ... 7
- Supporting figure S3. HERD-2.2 fiber diameter. ... 8
- Supporting figure S4. Additional TEM images of HERD-2.2 fibers with molecular crowding. ... 8
- Supporting figure S5. Additional TEM images of HERD-2.2 fibers in *E. coli*. ... 8
- Supporting figure S6. Additional TEM images of HERD-2.2–GFP condensates in *E. coli* ... 9
- Supporting figure S7. TEM images of wild type (WT) *E. coli*. ... 9
- Supporting figure S8. Confocal microscopy images of HERD-2.2 and HERD-2.2–GFP. ... 9
- Supporting figure S9. CLEM images of HERD-2.2 and HERD-2.2–GFP. ... 10
- Supporting figure S10. Confocal microscopy of HERD-2.2 fibers and fluorescent proteins. ... 11
- Supporting figure S11. HERD-2.2 and HERD-2.2–GFP *in vitro*. ... 12
- Supporting figure S12. Deconvolution of HERD-2.2 CD spectra. ... 13
- Supporting figure S13. TEM images of HERD-2.2 before and after heating to 90 °C ... 14
- Supporting figure S14. Image of HERD-2.2 fiber measured by X-ray fiber diffraction ... 14
- Supporting figure S15. X-ray fiber diffraction of HERD-2.2. ... 15
- Supporting figure S16. Change in transmission over time following cleavage. ... 16
- Supporting figure S17. SDS-PAGE of HERD-2.2-T–GFP cleavage ... 16
- Supporting figure S18. Confocal microscopy of HERD-2.2-T–GFP pre- and post-cleavage. ... 17
- Supporting figure S19. TEM images of HERD-2.2-T–GFP with and without added protease. ... 17

Supporting references ... 18

### Materials and methods

##### Materials

Lysogeny broth (LB), 2-amino-2-hydroxymethyl-1,3-propanediol (Tris), agar, phosphate buffered saline (PBS), ampicillin (AMP), chloramphenicol (CMP), isopropyl β-D-1-thiogalactopyranoside (IPTG), paraformaldehyde (PFA), imidazole, polyethylene glycol (PEG) 3350, urea, thrombin protease, thioflavin T, and sodium chloride (NaCl) were purchased from Sigma Aldrich. HisTrap HP immobilized metal ion affinity chromatography (IMAC), and HiLoad 16/600 Superdex size exclusion chromatography (SEC) columns were purchased from Cytiva. ProLong Diamond Antifade mountant was purchased from Invitrogen. cOmplete protease inhibitor tablets were purchased from Roche. Spin concentrator columns were purchased from Merck.

##### Computational tools

Protein parameters, including molecular weight and extinction coefficient (ϵ) at 280 nm were calculated from their primary amino acid sequences using ExPASy ProtParam (https://web.expasy.org/protparam).[1] All data analyses except those mentioned explicitly were performed in Python using the Pandas and Numpy libraries, and visualized using the MatPlotLib and Seaborn libraries.

##### Protein expression and purification

50 µl of chemically competent BL21*(DE3) *Escherichia coli* (*E. coli*) were transformed with the plasmid of interest by incubating on ice with DNA (typically 25 ng for a single vector, 100 ng each for two vectors) for 30 minutes, before heat shocking for 30 s at 42 °C, and chilling on ice for 1 minute. 200 µl of LB was then added and the cells incubated at 37 °C for 1 hour, before 50 µl of the cell suspension was spread on LB-agar plates containing appropriate antibiotics (AMP – 100 µg/ml, CMP – 25 µg/ml) and incubated at 37 °C overnight. For expression, a single colony was inoculated into a 5 ml LB supplemented with AMP or CMP, and grown overnight (37 °C, 200 rpm). Expression cultures (50 ml – 1 l) supplemented with AMP or CMP were inoculated 1:100 with overnight culture and grown to $OD_{600}$ = 0.4 – 0.6 (37 °C, 200 rpm). Protein expression was then induced to a final concentration of 400 µM IPTG and the culture grown for up to 21 hours (18 °C, 200 rpm).

For protein purification, cells were collected by centrifugation (3400 xg, 20 minutes) and resuspended in 40 ml resuspension buffer (20 mM Tris pH 7.5, 50 mM imidazole, 500 mM NaCl, 2 M urea, 1 tablet cOmplete protease inhibitor cocktail). Cells were lysed on ice by sonication (5 s on, 2 s off, 75% amplitude, 15 minutes). The suspension was centrifuged (16000 xg, 20 minutes) and filtered to clarify. The lysate was applied to a 5 ml HisTrap HP IMAC column pre-equilibrated in binding buffer (20 mM Tris pH 7.5, 500 mM NaCl, 50 mM imidazole, 2 M urea). The column was washed in binding buffer for 4 column volumes, and the bound protein eluted using a gradient of elution buffer (0 – 100%) (20 mM Tris pH 7.5, 500 mM NaCl, 2 M urea, 500 mM imidazole). The eluted protein was further purified by size exclusion chromatography using a HiLoad Superdex 200 pg column at 1 ml/min flow rate (20 mM Tris pH 7.5, 2 M urea). The purified protein was then buffer exchanged into 20 mM Tris pH 7.5 using a 26/10 desalting column run at 5 ml/min, before concentration using 3 kDa molecular weight cut-off (MWCO) spin concentrators, and snap frozen in liquid nitrogen for storage at -70 °C. Protein concentration was measured by absorbance at 280 nm.

##### Confocal microscopy

Microscopy was performed using a Leica SP8 confocal microscope using a 65 mW Ar laser (488 nm) with a 63x 1.4 numerical aperture oil immersion objective lens. For *in vitro* microscopy, 10 µl of sample was applied to a clean glass slide and covered with a coverslip before imaging. For in-cell imaging, *E. coli* treated as described above for expression of recombinant proteins. Samples were collected 5 hours after induction of protein expression for confocal microscopy. 1 ml of culture was collected by centrifugation (3000 xg, 3 minutes) and washed 3 times with 1 ml PBS, before fixing with 1 ml 2% PFA for 15 minutes. Fixed cells were washed 3 more times in PBS, before resuspending in 50 µl PBS. 10 µl was then applied to a clean glass slide, with 1 drop ProLong Diamond Antifade mountant, and covered with a glass coverslip. Cell images were analyzed and assembled in FiJi, and are displayed as maximum projections.[2,3]

**Transmission electron microscopy (TEM)**

Negative stain TEM was performed using a 120 kV Tecnai 12 electron microscope. For purified protein samples, 10 µl was applied to a glow-discharged 300-mesh carbon grid coated with pioloform and incubated for 1 minute, then stained using a 2% uranyl acetate (UA) solution. Grids were incubated for 1 second in a 10 µl drop of UA, then incubated sample-side down into a second drop and incubated for 3 minutes. The grid was then blotted with filter paper to remove excess stain, and swept through a third UA drop, and blotted again. The grid was then swept through two drops of ultrapure water, before blotting and leaving to dry before imaging.

For TEM of *E. coli*, cultures treated as described above for expression of recombinant proteins were collected 21 hours after induction of protein expression. 1 ml of cell suspension was pelleted by centrifugation, and 1 µl of pellet was vitrified using a Leica EM PACT2 high pressure freezer with a rapid transfer system. The vitrified cells were freeze substituted with 0.2% UA, 5% $H_2O$ in acetone for 5 hours at -90 °C using a Leica AFS2 automated freeze substitution system. The samples were then warmed to -45 °C, and kept for 2 hours at that temperature before washing in acetone for 30 minutes. Resin (Lowicryl HM20) was then infiltrated into the freeze substituted samples at increasing dilutions (25%, 50%, 75%) for 3 hours each, before embedding in 100% resin for 16 hours, followed by 3 changes of resin, left for 2 hours each. The infiltrated resin was then polymerized using UV for 48 hours. The resin blocks were sectioned using an EM UC6 microtome with a diamond knife at 45 °, and the sections imaged by TEM.

**Correlative light electron microscopy (CLEM)**

Samples were prepared for CLEM as described for TEM, but sections were first imaged using a Leica SP8 AOBS confocal microscope with a 63x oil immersion objective, before negative staining with 2% UA as described for purified samples, and imaging by TEM.

**Circular dichroism (CD) spectroscopy**

Circular dichroism (CD) spectra were recorded on a a JASCO J-810 spectropolarimeter with a Peltier temperature controller, in a 10 mm path length reduced volume cuvette. Full spectra measured ellipticity between 190 nm and 260 nm in 1 nm intervals, with a 100 nm/min scanning rate, 1 nm bandwidth and 1 s response time. A reference spectrum using the same cuvette, parameters and buffer at 5 °C was subtracted from the measured ellipticity. Measurements of ellipticity with respect to temperature (melting and cooling spectra), were recorded by collecting initial spectra at 5 °C, with ellipticity measured at 222 nm every 1 °C and full spectra measured every 5 °C as the temperature was increased to 90 °C, and then decreased again to 5 °C. Ellipticity (deg) values were converted to mean residue ellipticity (MRE) ($deg \cdot cm^2 \cdot dmol^{-1} \cdot res^{-1}$) by normalization to the number of peptide bonds in the protein, and the path length using the following equation:

$$\boldsymbol{MRE\ (deg \cdot cm^2 \cdot dmol^{-1} \cdot res^{-1}) = \frac{\theta \times 100}{c \times l \times b}}$$

Where $\theta$ is the difference in absorbed circularly polarized light in millidegrees, $c$ is the protein concentration in mM, $l$ is the path length in cm, and $b$ is the number of amide bonds in the protein. Fraction helicity was calculated using the MRE at 222 nm ($MRE_{222}$) using the following equation:

$$\textbf{\textit{Fraction helix}}\ (\%) = \mathbf{100} \times \frac{\boldsymbol{MRE_{222} - MRE_{coil}}}{\boldsymbol{-42500 \times \left(1 - \frac{3}{n}\right) - MRE_{coil}}}$$

Where $MRE_{coil}$ is 640 - 45T, T is the temperature in degrees Celsius, and $n$ is the number of amide bonds in the sample.[4] Melting temperature (Tm) was calculated as the mean of 3 independent experiments. Deconvolution of CD data was performed using https://bestsel.elte.hu/index.php.[5]

**X-ray fiber diffraction**

Solutions containing purified fibers (16 mg/ml protein, 20 mM Tris pH 7.5) were hung between two glass capillaries with the ends sealed with paraffin, placed approximately 1 cm apart, and left to dry for 18 hours. Fibers were aligned in the detector at 0 and 90° orientations, before diffraction using a Rigaku copper rotating anode X-ray source with a Saturn CCD detector at distances of 50 and 100 mm for 30 – 60 s. Diffraction patterns were analyzed using CLEARER to identify distances.[6]

**Thioflavin T fluorescence**

Thioflavin T (ThT) fluorescence (excitation: 449 nm, emission: 493 nm) was measured using a CLARIOstar plate reader in black 96-well plates. Solutions containing 25 µM ThT, 50 mM Tris pH 7.5, and varied protein concentration from 0 – 100 µM were made up to 100 µl per well, with 4 replicates per protein concentration. Plates were sealed to prevent evaporation, and incubated at 25 °C for 5 hours to equilibrate, before measuring fluorescence emission every 85 seconds for 12 minutes. The mean fluorescence over this period was recorded as the final fluorescence intensity for each well, and plotted as mean and standard deviation over the 4 replicates.

**Protease cleavage and droplet maturation**

Droplet maturation was triggered by addition of thrombin protease to phase-separated droplets. Phase-separated droplets were formed by mixtures of 10% PEG 3350, 125 mM NaCl, 50 mM Tris pH 7.5, and 0.675 mM HERD-2.2-T–GFP, before 0.06 units of thrombin protease added, and incubated at 25 °C for 30 minutes. Complete cleavage was determined by SDS-PAGE after 1 hour.

#### Supporting information

| **Protein name** | **Sequence** |
|---|---|
| HERD-2.2 | MGSHHHHHHHHHHENLYFQSGSGSGTMGSGIKEEAAAIKWEAAAIKEGASP<br>EPQPKPSGDPQSKQTPEPSRSQGQAKEIQWQAKEIQEQAKG* |
| HERD-2.2–GFP | MGSHHHHHHHHHHENLYFQSGSGSGTMGSGIKEEAAAIKWEAAAIKEGASP<br>EPQPKPSGDPQSKQTPEPSRSQGQAKEIQWQAKEIQEQAKGGSGSGHHMVS<br>KGEELFTGVVPILVELDGDVNGHKFSVSGEGEGDATYGKLTLKFICTTGKLPV<br>PWPTLVTTLTYGVQCFARYPDHMKQHDFFKSAMPEGYVQERTIFFKDDGNY<br>KTRAEVKFEGDTLVNRIELKGIDFKEDGNILGHKLEYNYNSHKVYITADKQK<br>NGIKVNFKTRHNIEDGSVQLADHYQQNTPIGDGPVLLPDNHYLSTQSKLSKD<br>PNEKRDHMVLLEFVTAAGITLGMDELYK* |
| HERD-2.2–mCherry | MGSHHHHHHHHHHENLYFQSGSGSGTMGSGIKEEAAAIKWEAAAIKEGASP<br>EPQPKPSGDPQSKQTPEPSRSQGQAKEIQWQAKEIQEQAKGGSGSGHHMVS<br>KGEEDNMAIIKEFMRFKVHMEGSVNGHEFEIEGEGEGRPYEGTQTAKLKVT<br>KGGPLPFAWDILSPQFMYGSKAYVKHPADIPDYLKLSFPEGFKWERVMNFED<br>GGVVTVTQDSSLQDGEFIYKVKLRGTNFPSDGPVMQKKTMGWEASSERMYP<br>EDGALKGEIKQRLKLKDGGHYDAEVKTTYKAKKPVQLPGAYNVNIKLDITSH<br>NEDYTIVEQYERAEGRHSTGGMDELYK* |
| HERD-2.2-T–GFP | MGSHHHHHHHHHHENLYFQSGSGSGTMGSGIKEEAAAIKWEAAAIKEGASP<br>EPQPKPSGDPQSKQTPEPSRSQGQAKEIQWQAKEIQEQAKGGLVPRGSSGSG<br>HHMVSKGEELFTGVVPILVELDGDVNGHKFSVSGEGEGDATYGKLTLKFICTT<br>GKLPVPWPTLVTTLTYGVQCFARYPDHMKQHDFFKSAMPEGYVQERTIFFK<br>DDGNYKTRAEVKFEGDTLVNRIELKGIDFKEDGNILGHKLEYNYNSHKVYIT<br>ADKQKNGIKVNFKTRHNIEDGSVQLADHYQQNTPIGDGPVLLPDNHYLSTQS<br>KLSKDPNEKRDHMVLLEFVTAAGITLGMDELYK* |
| GFP | MVSKGEELFTGVVPILVELDGDVNGHKFSVSGEGEGDATYGKLTLKFICTTGK<br>LPVPWPTLVTTLTYGVQCFARYPDHMKQHDFFKSAMPEGYVQERTIFFKDD<br>GNYKTRAEVKFEGDTLVNRIELKGIDFKEDGNILGHKLEYNYNSHKVYITADK<br>QKNGIKVNFKTRHNIEDGSVQLADHYQQNTPIGDGPVLLPDNHYLSTQSKLS<br>KDPNEKRDHMVLLEFVTAAGITLGMDELYK* |
| mCherry | MVSKGEEDNMAIIKEFMRFKVHMEGSVNGHEFEIEGEGEGRPYEGTQTAKL<br>KVTKGGPLPFAWDILSPQFMYGSKAYVKHPADIPDYLKLSFPEGFKWERVMN<br>FEDGGVVTVTQDSSLQDGEFIYKVKLRGTNFPSDGPVMQKKTMGWEASSER<br>MYPEDGALKGEIKQRLKLKDGGHYDAEVKTTYKAKKPVQLPGAYNVNIKLDI<br>TSHNEDYTIVEQYERAEGRHSTGGMDELYK* |
| CC-Di* | Ac-GEIAALKQEIAALKKENAALKWEIAALKQG-$NH_2$ |

**Supporting table S1.** Protein sequences used in this study.

*CC-Di was synthesized by solid-phase peptide synthesis as a *C*-terminal amide and *N*-terminally acetylated peptide.

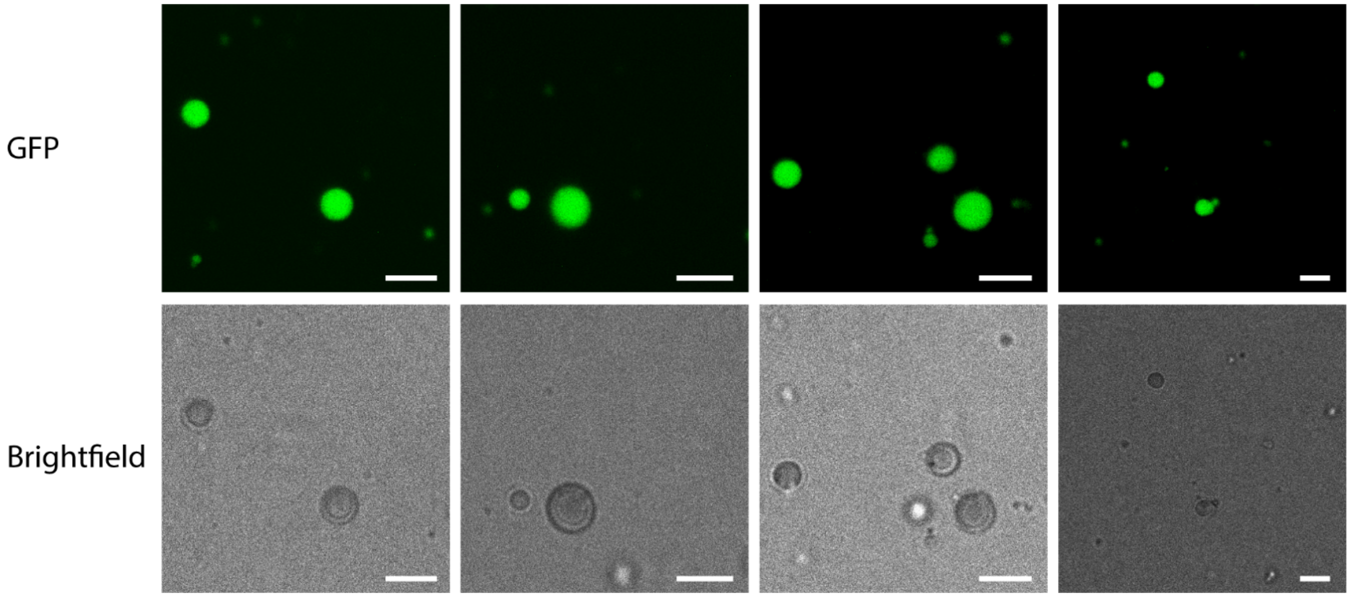


**Supporting figure S1.** Additional confocal microscopy images of HERD-2.2–GFP droplets. Conditions: 110 µM HERD-2.2–GFP, 10% PEG 3350, 125 mM NaCl, 50 mM Tris pH 7.5. Scale bars 5 µm.

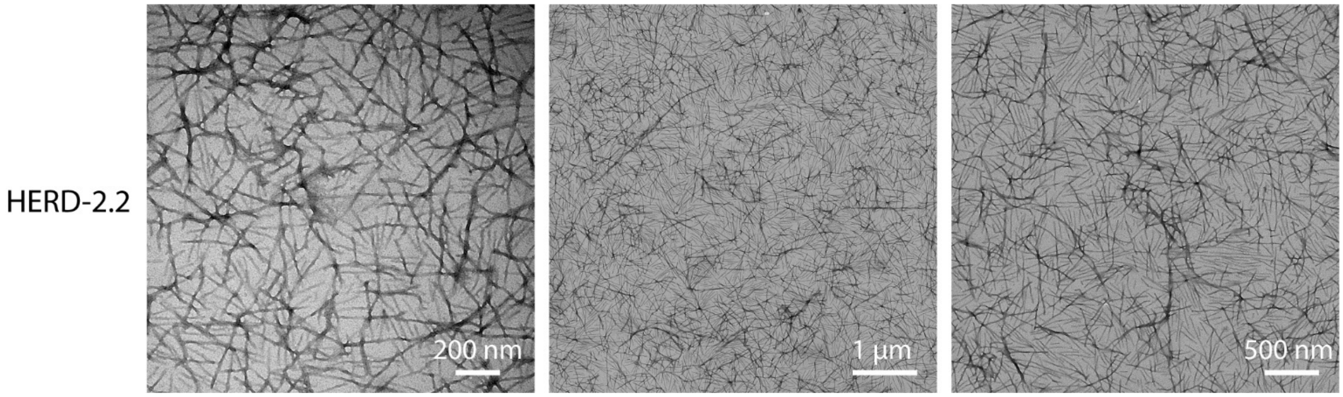


**Supporting figure S2.** Additional TEM images of HERD-2.2 fibers. Conditions: 25 µM HERD-2.2, 50 mM Tris pH 7.5.

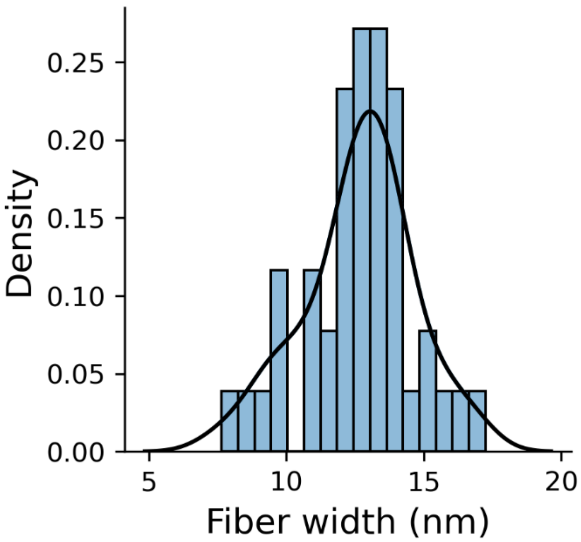


**Supporting figure S3.** HERD-2.2 fiber diameter.

Diameter measured from 43 HERD-2.2 fibers without molecular crowding agents. Kernel density estimation (KDE) computed from the histogram is plotted as a solid black line.

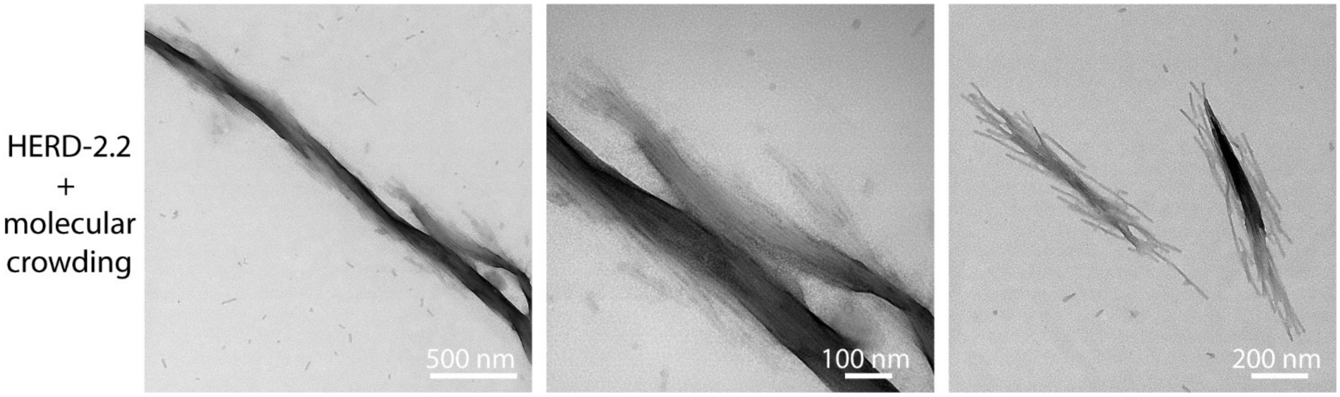


**Supporting figure S4.** Additional TEM images of HERD-2.2 fibers with molecular crowding.

Conditions: 25 µM HERD-2.2, 10% PEG 3350, 125 mM NaCl, 50 mM Tris pH 7.5.

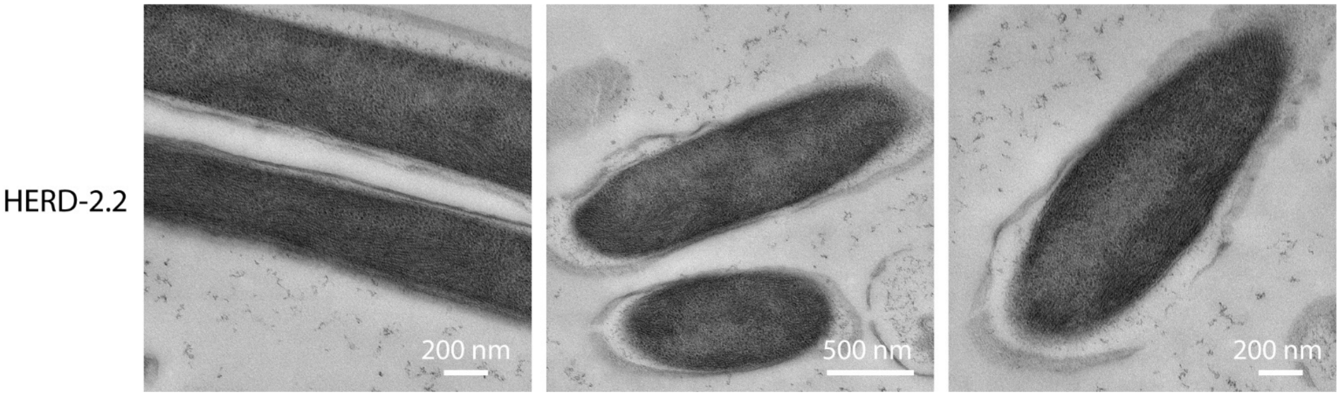


**Supporting figure S5.** Additional TEM images of HERD-2.2 fibers in *E. coli*.

Cells collected and vitrified 21 hours after induction of expression.

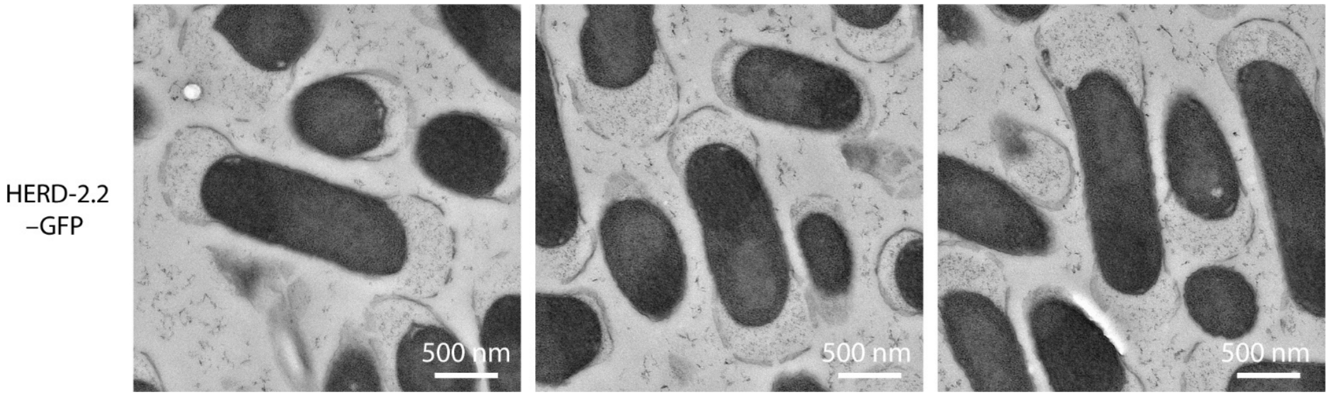


**Supporting figure S6.** Additional TEM images of HERD-2.2–GFP condensates in *E. coli*.
Cells collected and vitrified 21 hours after induction of expression.

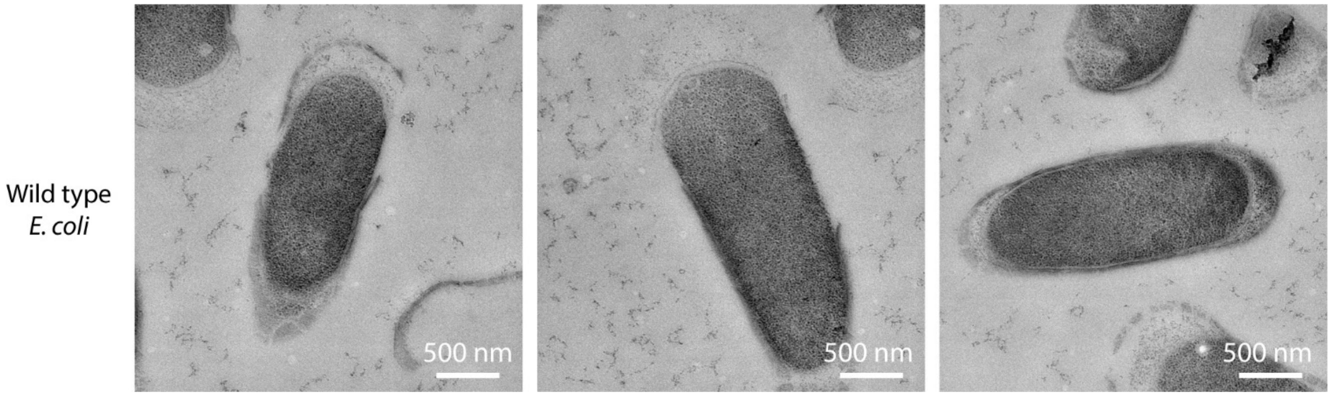


**Supporting figure S7.** TEM images of wild type (WT) *E. coli*.
Cells collected and vitrified 21 hours after induction of expression.

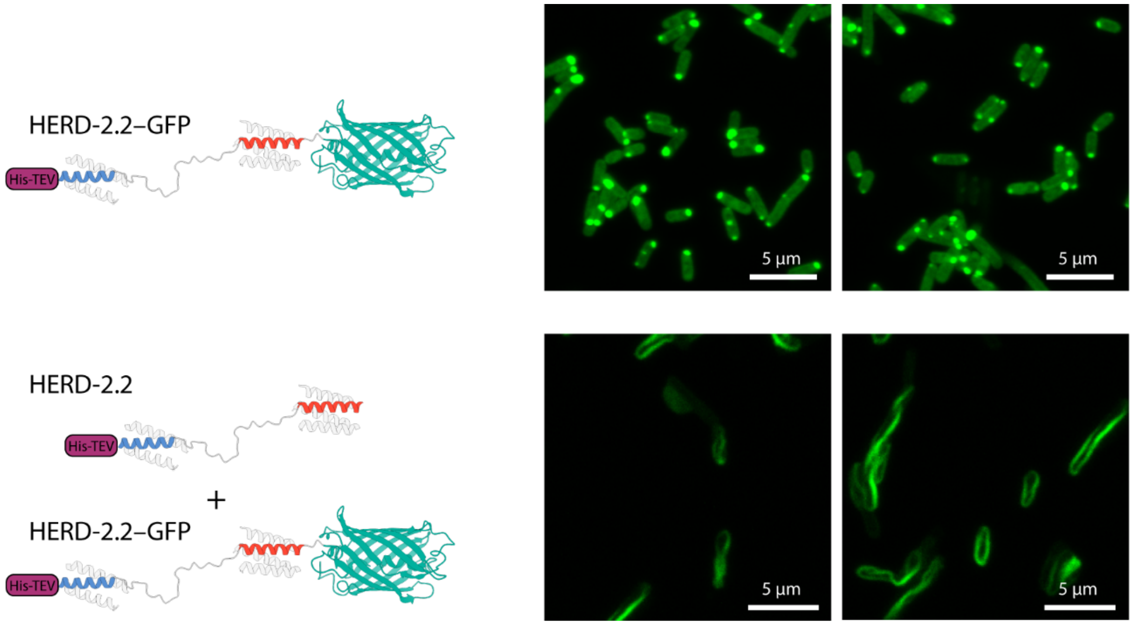


**Supporting figure S8.** Confocal microscopy images of HERD-2.2 and HERD-2.2–GFP.
Cells collected and fixed 5 hours after induction of expression.

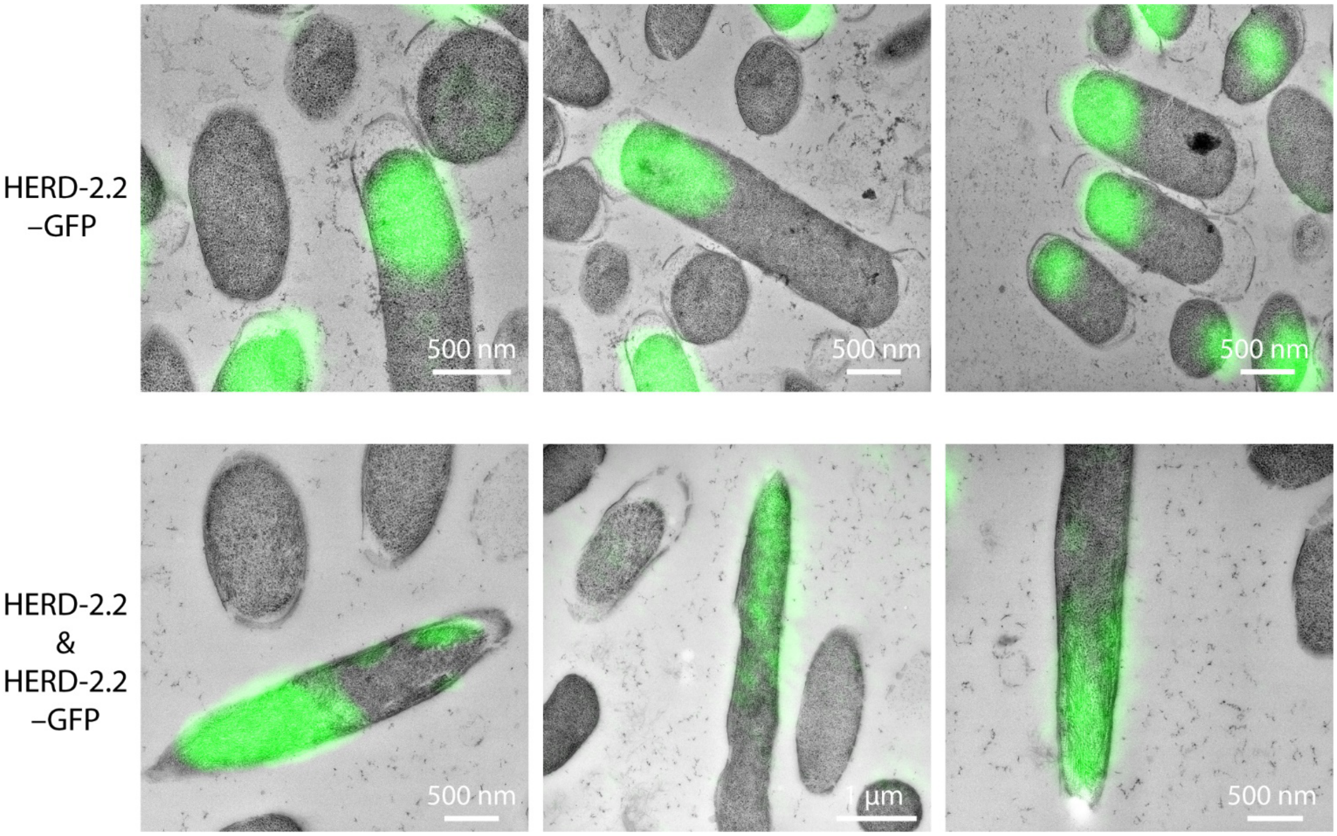


**Supporting figure S9.** CLEM images of HERD-2.2 and HERD-2.2–GFP.
Cells collected and vitrified 21 hours after induction of expression.

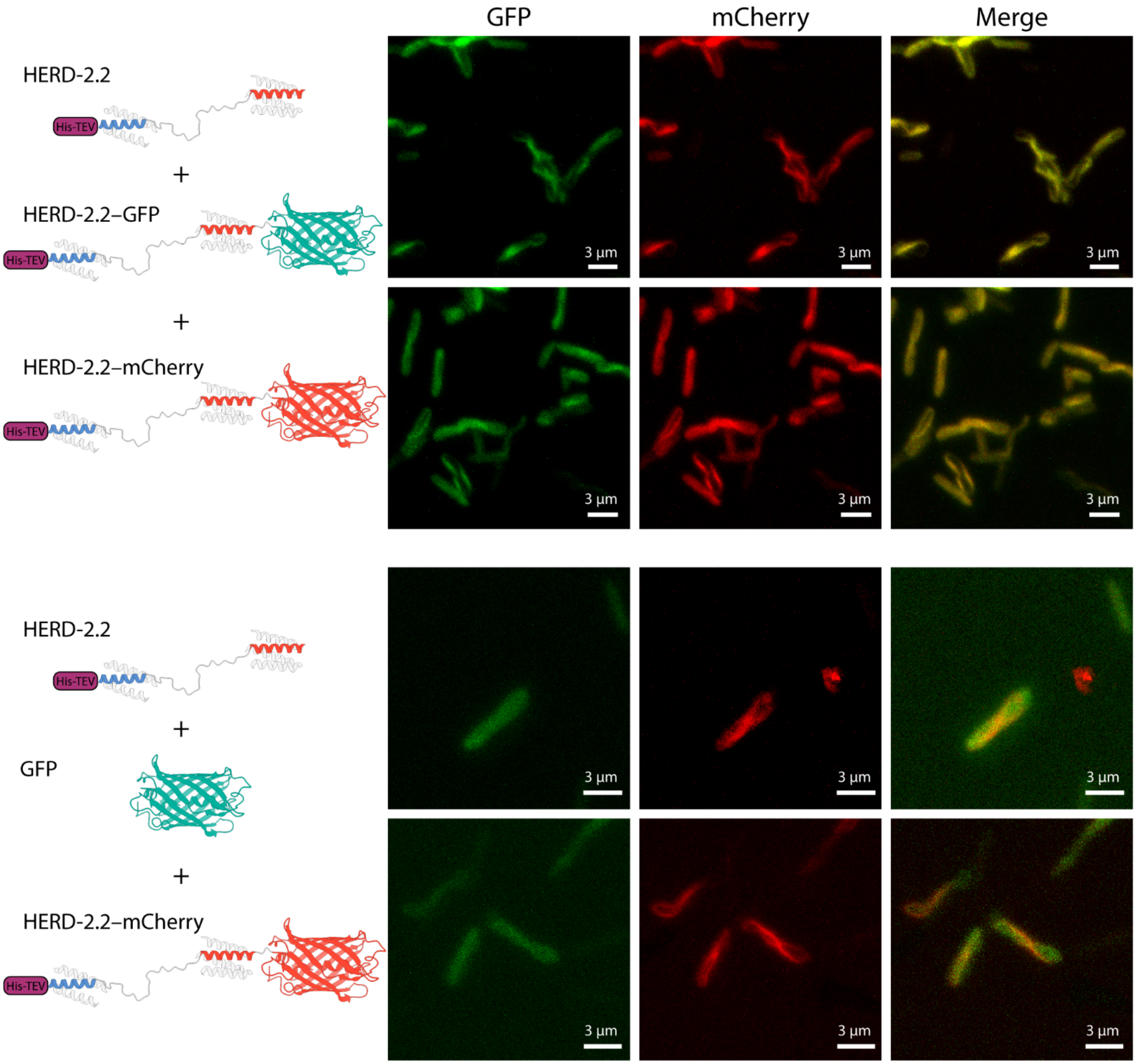


**Supporting figure S10.** Confocal microscopy of HERD-2.2 fibers and fluorescent proteins. Cells collected and fixed 5 hours after induction of expression.

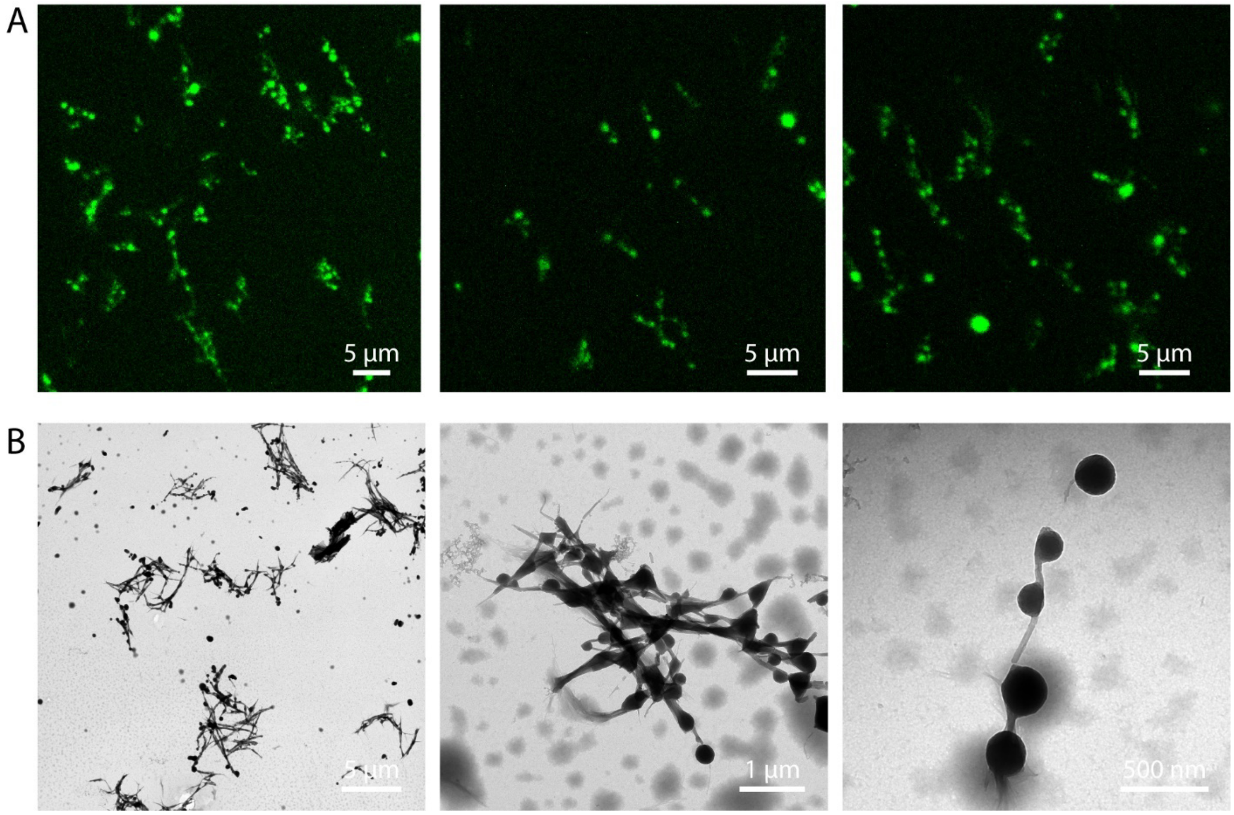


**Supporting figure S11.** HERD-2.2 and HERD-2.2–GFP *in vitro*.

Confocal microscopy (A) and TEM (B) of HERD-2.2 and HERD-2.2–GFP mixtures *in vitro*. Conditions: (A) 82 µM HERD-2.2, 110 µM HERD-2.2–GFP, 10% PEG 3350, 125 mM NaCl, 50 mM Tris pH 7.5. (B) 82 µM HERD-2.2, 55 µM HERD-2.2–GFP, 10% PEG 3350, 125 mM NaCl, 50 mM Tris pH 7.5.

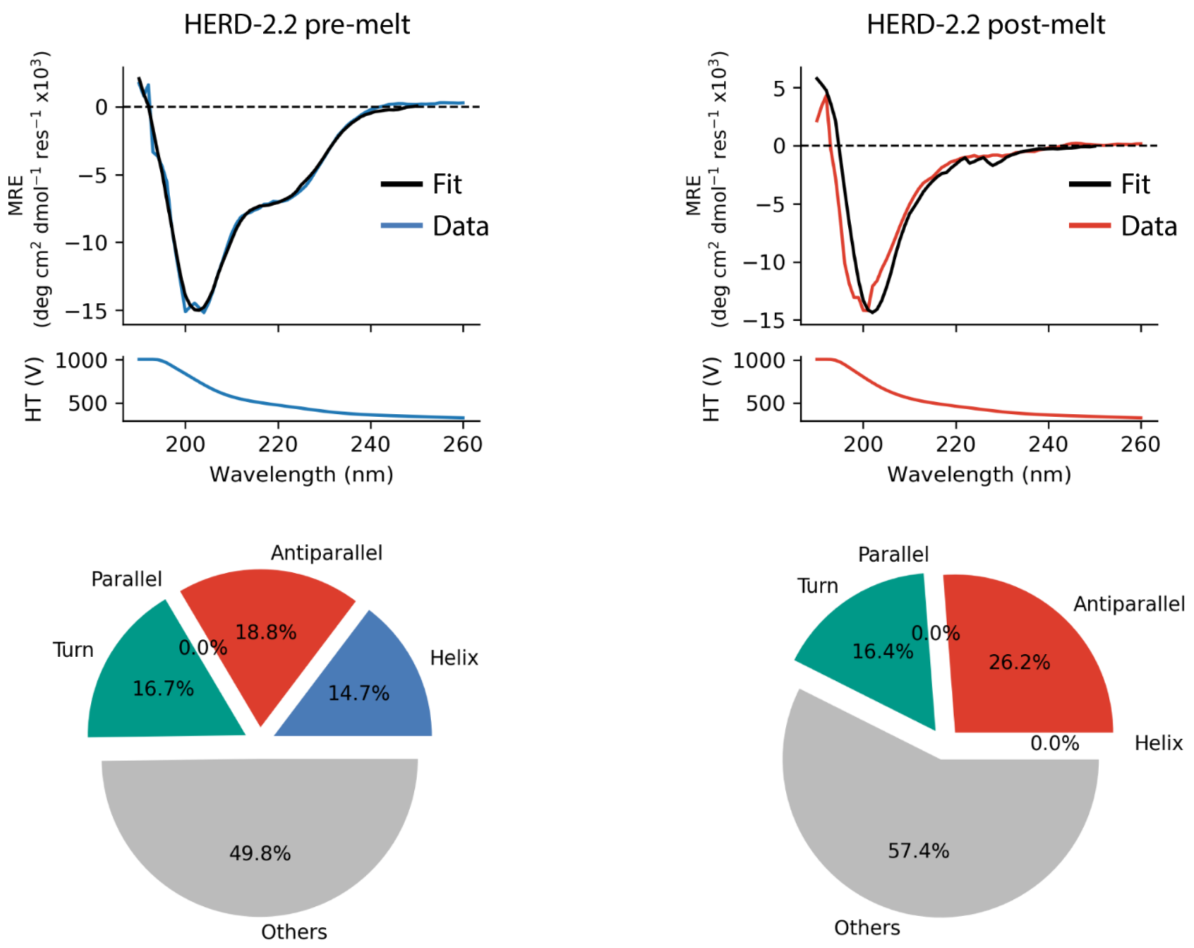


**Supporting figure S12.** Deconvolution of HERD-2.2 CD spectra.

Deconvolution was performed with https://bestsel.elte.hu/index.php.[5] CD conditions: 25 μM HERD-2.2, 50 mM Tris pH 7.5.

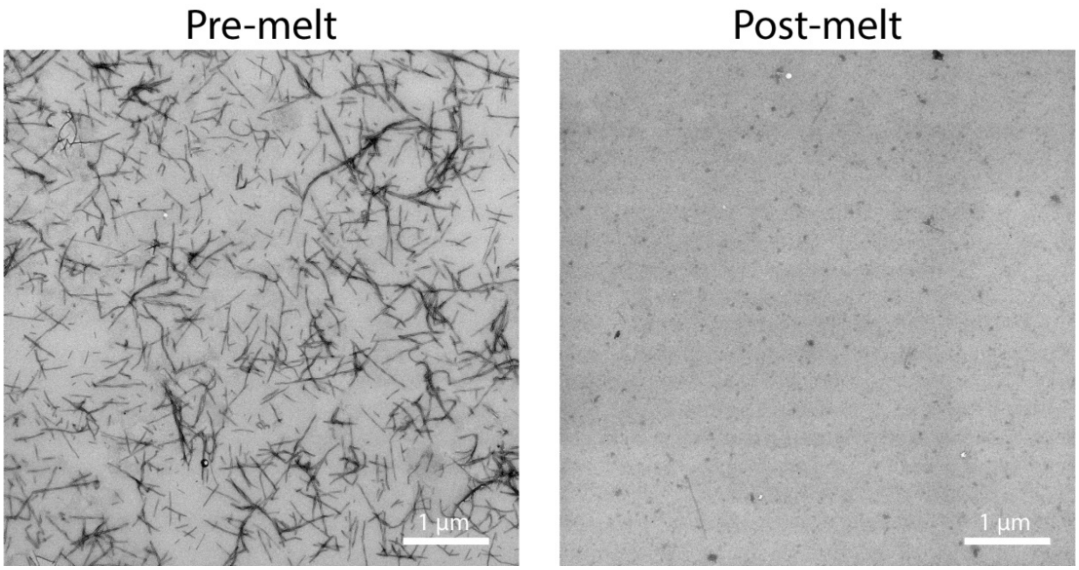


**Supporting figure S13.** TEM images of HERD-2.2 before and after heating to 90 °C. Conditions: 25 µM HERD-2.2, 50 mM Tris pH 7.5.

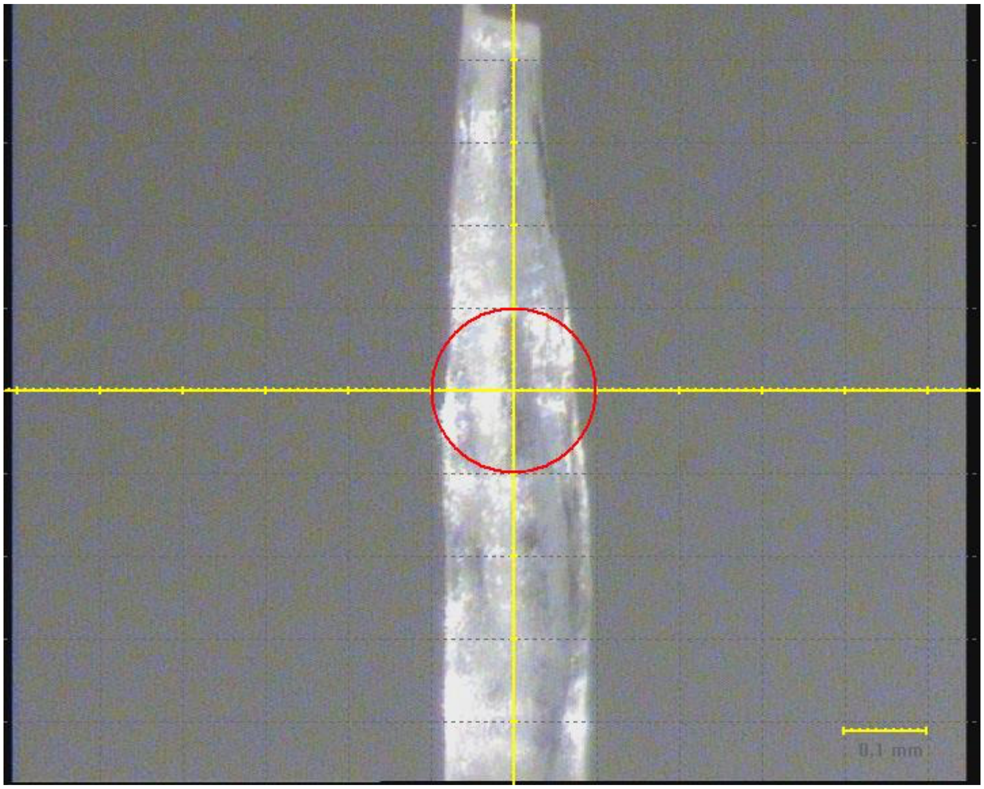


**Supporting figure S14.** Image of HERD-2.2 fiber measured by X-ray fiber diffraction. Conditions: 1.6 mM HERD-2.2, 50 mM Tris pH 7.5.

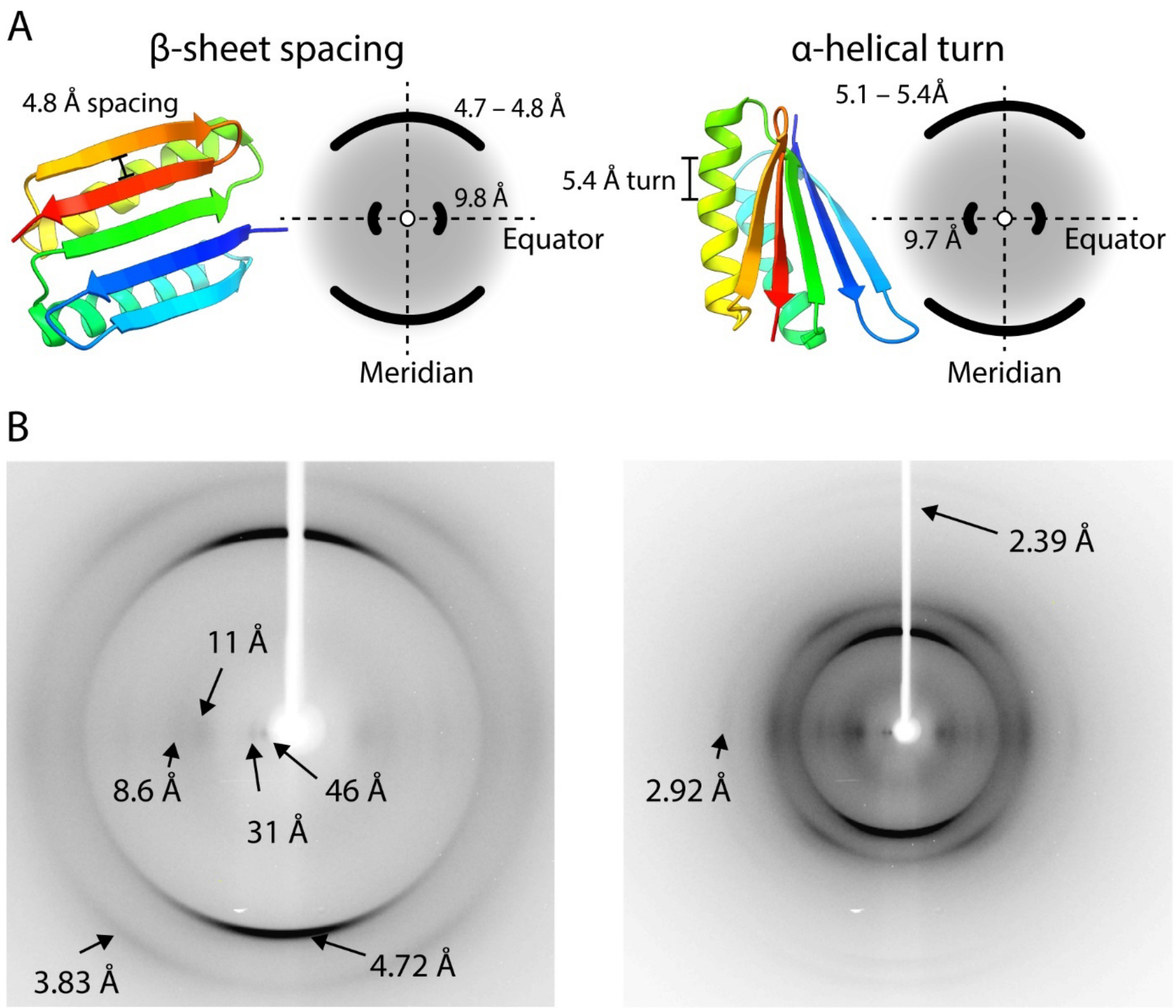


**Supporting figure S15.** X-ray fiber diffraction of HERD-2.2.

(A) Cartoon representation of the expected major equatorial and meridional reflections produced by α-helical and cross-β fibrils. (B) Fiber diffraction patterns for HERD-2.2 collected at 100 mm (left) and 50 mm (right). Conditions: 1.6 mM HERD-2.2, 50 mM Tris pH 7.

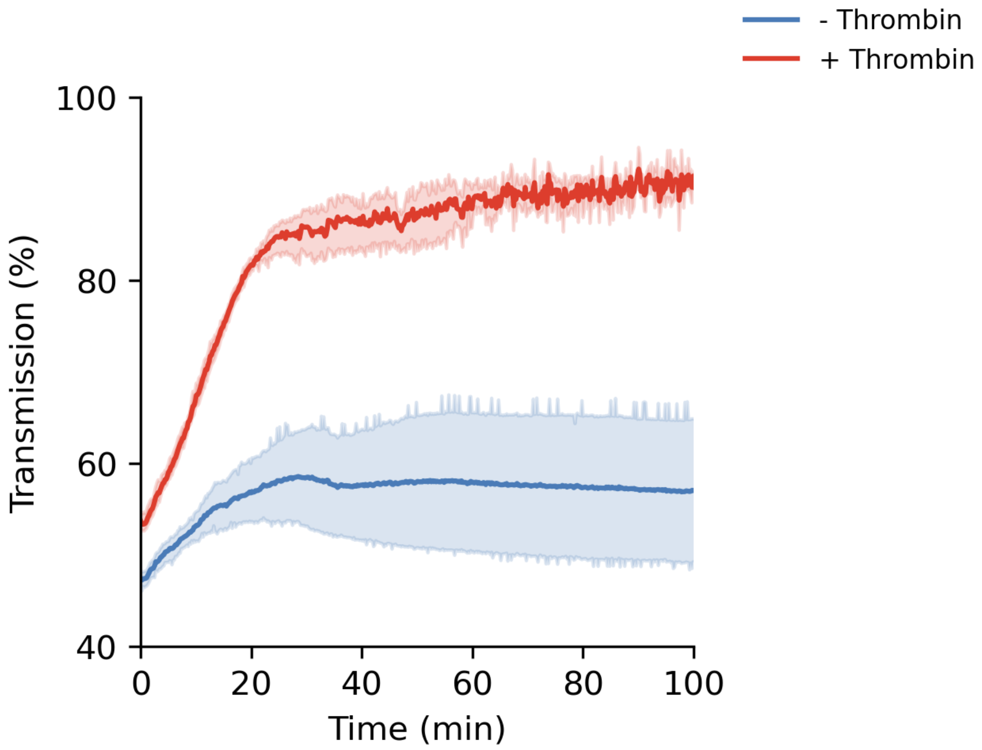


**Supporting figure S16.** Change in transmission over time following cleavage.

Time 0 is the point of phase separation, and addition of protease. From n = 4 independent repeats. Conditions: 10% PEG 3350, 125 mM NaCl, 50 mM Tris pH 7.5, 0.675 mM HERD-2.2-T–GFP, in a total volume of 80 µl. + Thrombin had additional 0.06 units of thrombin protease added.

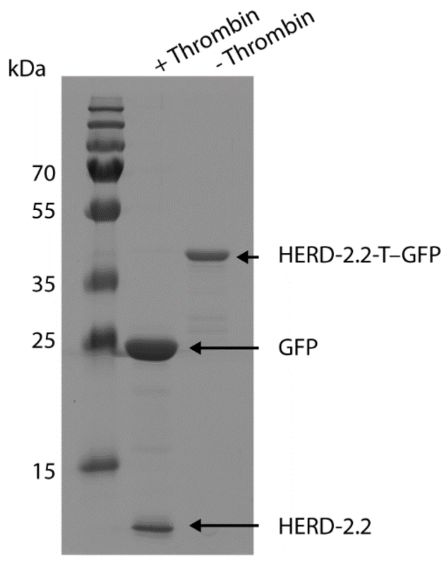


**Supporting figure S17.** SDS-PAGE of HERD-2.2-T–GFP cleavage.

17% acrylamide/bis-acrylamide SDS-PAGE gel of HERD-2.2-T–GFP with (+thrombin) and without (– thrombin) addition of 0.06 units of thrombin protease.

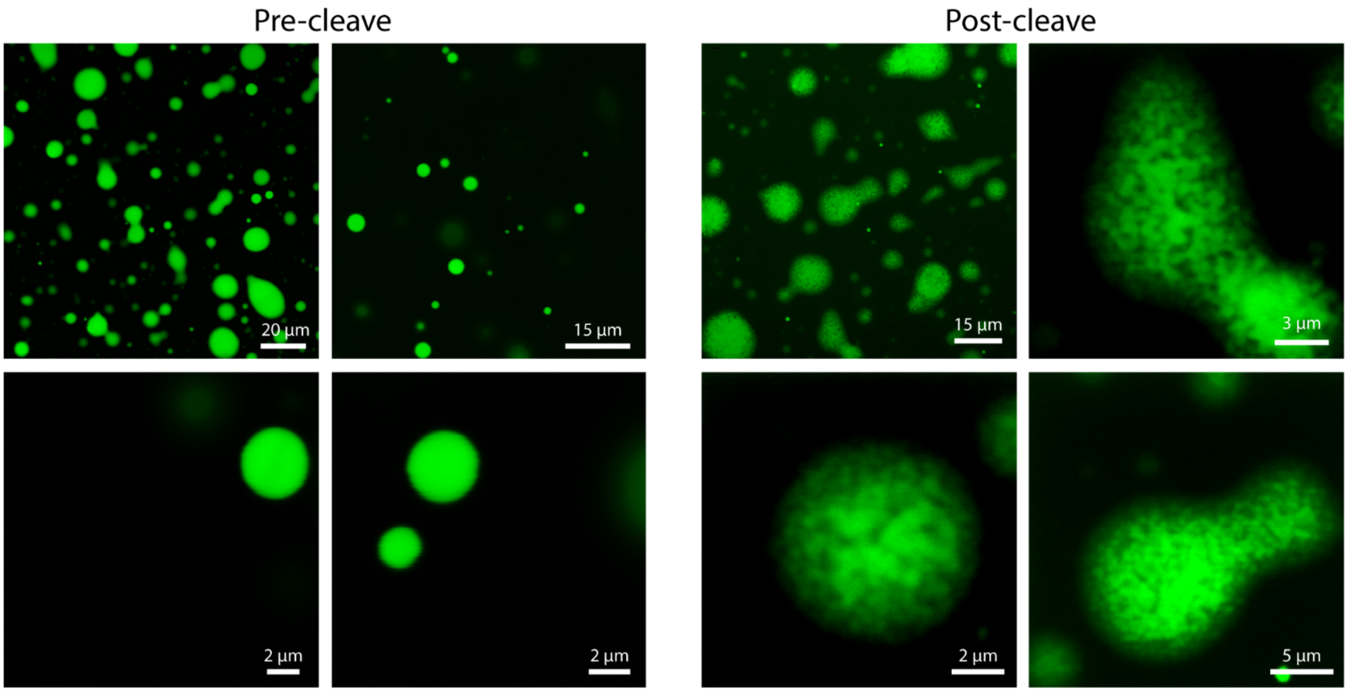


**Supporting figure S18.** Confocal microscopy of HERD-2.2-T–GFP pre- and post-cleavage.

Post-cleave was 30 minutes after addition of thrombin. Conditions: 10% PEG 3350, 125 mM NaCl, 50 mM Tris pH 7.5, 0.675 mM HERD-2.2-T–GFP with (+ thrombin) or without (– thrombin) addition of 0.06 units of thrombin protease.

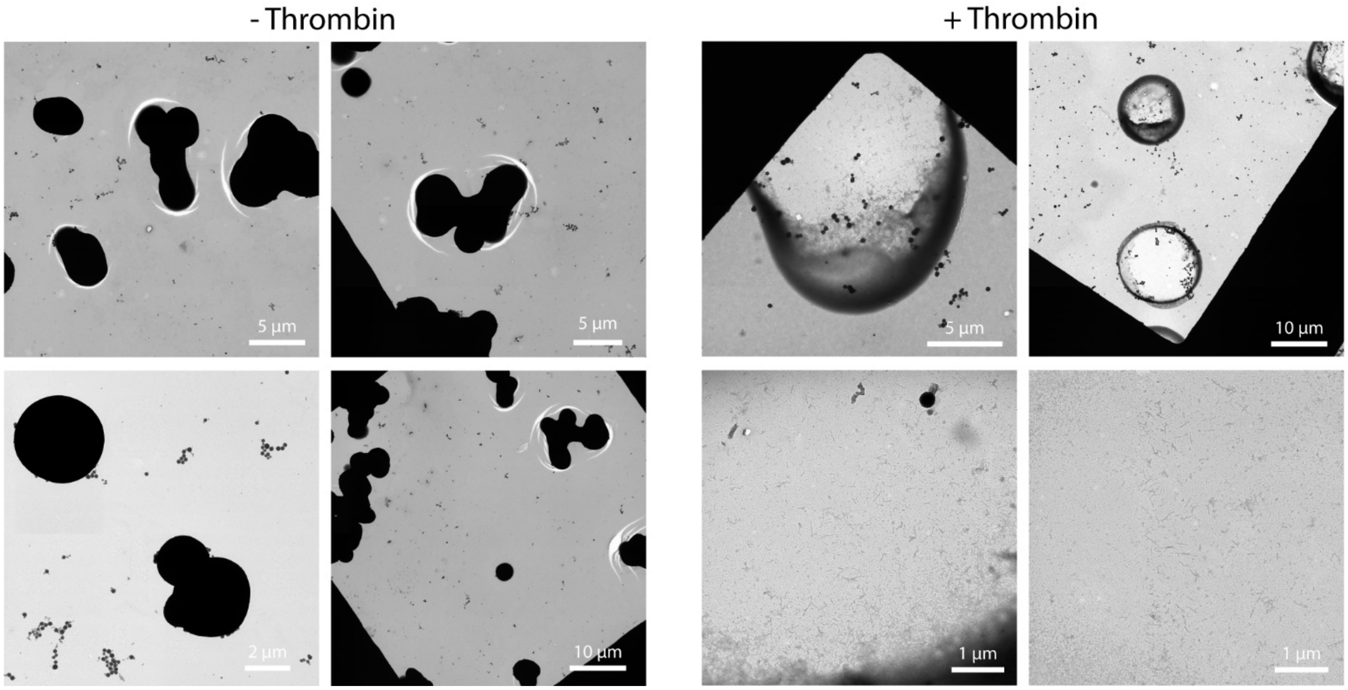


**Supporting figure S19.** TEM images of HERD-2.2-T–GFP with and without added protease.

Both samples were imaged after 30 minutes. Conditions: 10% PEG 3350, 125 mM NaCl, 50 mM Tris pH 7.5, 0.675 mM HERD-2.2-T–GFP with (+ thrombin) or without (– thrombin) addition of 0.06 units of thrombin protease.
